# Ontogeny of the organized nasopharynx-associated lymphoid tissue in rainbow trout

**DOI:** 10.1101/2023.08.04.552019

**Authors:** Benjamin J Garcia, Alexis Reyes, Chrysler Martinez, Yago Serra dos Santos, Irene Salinas

**Author notes:** Corresponding author: Dr. Irene Salinas, Department of Biology, University of New Mexico, Albuquerque, NM87131, USA.

## Abstract

Understanding the ontogeny of teleost mucosa-associated lymphoid tissues (MALT) is critical for determining the earliest timepoint for effective mucosal vaccination of young fish. Here, we describe the developmental sequence that leads to the formation of an organized MALT structure in rainbow trout, the organized nasopharynx-associated lymphoid tissue (O-NALT). Control rainbow trout were sampled between 340 and 1860 degree days (DD) and routine histology and immunofluorescence staining were used to determine cellular changes in immune cells in the nasal cavity as well as O-NALT formation. We identified that O-NALT is first seeded by CD8α^+^ T cells at 700DD followed by IgM^+^ B cells and CD4-2b^+^ T cells at 1000DD. Histomorphologically, trout O-NALT is fully formed at 1400DD when it is composed of 67% CD4-2b^+^ cells, 20% IgM^+^ cells, 13% CD8α^+^ T cells, and no IgT^+^ B cells. Whole body gene expression analyses uncovered waves of *igmh*, *cd4-2b*, and *cd8a* expression that recapitulate the cellular seeding sequence of O-NALT by specific lymphocyte subsets. Our results indicate that 1) O-NALT formation results from a specific sequence of lymphocyte subset colonization pioneered by CD8α^+^ T cells and 2) the presence of the full O-NALT structure at 1400DD may mark this timepoint as the earliest developmental stage at which mucosal vaccines can induce long lasting, specific immune responses.

## 1. Introduction

Vaccine delivery in finfish must be closely timed to the development of the adaptive immune system since fish are at their most vulnerable during early life stages (Buchmann, 2022). The teleost adaptive immune system does not fully develop until well after hatching. For instance, in rainbow trout (*Oncorhynchus mykiss*), Ig^+^ cells can be detected as early as 4 days post-hatch (dph) in the head kidney but are only seen in the spleen at around 30 dph (Rombout et al., 2011; Solem and Stenvik, 2006; Zapata et al., 2006). In both spleen and head kidney, the timing of the appearance of Ig^+^ cells corresponds to the first appearance of high levels of MHC-I protein expression, potentially indicating a developmentally orchestrated cascade (Fischer et al., 2005). In relation to T cell mediated immunity, the thymus in rainbow trout develops as early as 8 days pre-hatch, but allograft responses are not observed until 200DD, indicating that early lymphocytes in the thymus are likely not functional until 200DD (Tatner, 1986; Zapata et al., 2006). Specific humoral immune responses to intraperitoneally-delivered antigens in rainbow trout have not been observed until 800DD, indicating that the functional development of the adaptive immune system occurs well after the presence of its individual component cell types (Tatner, 1986). Yet, in depth ontogeny studies of lymphoid organs, cell types and immune markers in rainbow trout are still missing.

In the teleost mucosa, the development of adaptive immunity lags well behind that of systemic lymphoid organs (Salinas et al., 2011). For example, common carp IgM^+^ B cells are first detected in whole body samples around 2 weeks post-fertilization (wpf), with the first plasma cells present around 4 wpf. However, the first IgM^+^ cells do not appear in the intestine and gills of this species until 6-7 wpf (Romano et al., 1997). Another study in carp found that gene expression of IgT/IgZ, an Ig isotype specialized in mucosal immunity, appears in the gut later during ontogeny than IgM, further cementing the view that mucosal immunity lags systemic immunity (Ryo et al., 2010). Similarly, seeding of B and T cells at mucosal tissues lags the presence of these cell types in spleen and other lymphoid tissues in sea bass (Picchietti et al., 1997) and spotted wolf fish (Grontvedt and Espelid, 2003).

The mucosa-associated lymphoid tissue (MALT) in teleosts has traditionally been described as a diffuse network of immune cells scattered intraepithelially and in the lamina propria of the mucosal epithelium (Miller and Salinas, 2020; Salinas et al., 2011). Challenging this dogma, several studies have identified lymphoid structures associated with several mucosal barriers in teleosts including the interbranchial lymphoid tissue (ILT) (Aas et al., 2017; Dalum et al., 2015; Koppang et al., 2010) the amphibranchial lymphoid tissue (ALT) (Dalum et al., 2021), the bursa (Loken et al., 2020) and the Neumausean lymphoid organ (NEMO) (Resseguier et al., 2023). A rough ontogeny of the ILT has been determined in Atlantic salmon (*Salmo salar*): the ILT appears somewhere between the yolk-sac and juvenile stages at a body weight between 0.22g and 36.3g, but the exact time point of appearance, or specific events that lead to the formation of this structure have not been determined (Dalum et al., 2016). Importantly, the developmental time at which the ILT becomes functional is still unknown. While no specific B and T cells stains have outlined the ontogeny of NEMO, observations based on publicly available H&E stains of zebrafish sections indicate that a structure reminiscent of the NEMO appears between 4 and 6 weeks of development, around the same time that the zebrafish adaptive immune system is considered fully developed (Resseguier et al., 2023). Finally, the anatomical and functional ontogeny of the ALT and the bursa have not yet been determined. Thus, there is a knowledge gap in our understanding of teleost MALT ontogeny especially when it comes newly discovered organized or semi-organized MALT.

Recently, our group described an organized nasopharynx associated lymphoid tissue (O-NALT) in rainbow trout composed of 24% IgM^+^ B cells, 56% CD4-2b^+^ T cells, 16% CD8α^+^ T cells, and 4% IgT^+^ B cells at the 30g stage. Trout O-NALT expresses molecular markers of mammalian germinal centers (Garcia et al., 2022) suggesting its critical role in adaptive immune responses. Following intranasal vaccination, B cells proliferate and enter apoptosis in trout O-NALT and *aicda* expression (the gene encoding for AID, the enzyme that mediates somatic hypermutation) increases significantly relative to controls. This is in sharp contrast with responses in the olfactory rosette, where no organized lymphoid structures have been found, and B cell proliferation and apoptosis do not change significantly in vaccinated trout (Garcia et al., 2022). Our work on O-NALT was performed in juvenile animals weighing around 30g, and the developmental changes that occur until this structure is formed are currently unknown.

Proper timing of mucosal vaccination in young animals requires an understanding of MALT ontogeny. Our previous work showed that intranasal administration of a formalin-killed vaccine in rainbow trout against enteric red mouth (ERM) disease as early as 360DD can provide some protection against *Yersinia ruckeri* challenge, which lasts for at least 28 days following vaccination (Salinas et al., 2015). This contrasts with intranasal administration of a live-attenuated viral vaccine against infectious hematopoietic necrosis virus (IHNV) at the same age, which resulted in 40% mortality, a result not seen in animals which received the same vaccine intranasally at 450DD. In the same study, we captured histological changes in the olfactory organ of rainbow trout from 360DD to 1050DD. However, because at the time we did not know of the existence of O-NALT, all our data focused on the presence of scattered leucocytes in the olfactory organ and not the organized lymphoid structure found in association with the wall of the nasal cavity (O-NALT). Thus, the main goal of this study is to understand the sequence of events that lead to the formation of O-NALT in unvaccinated rainbow trout using cellular and molecular markers to establish optimal vaccination windows for intranasal delivery.

## 2. Methods

### 2.1. Husbandry

Juvenile rainbow trout (*Oncorhynchus mykiss*) (body weight approximately 0.1g) were obtained from Trout Lodge (Bonney Lake, WA) at 340DD of age, and were housed in the University of New Mexico aquatic animal housing facility. Animals were maintained at a 12-h dark/12-h light photoperiod and fed a commercial pellet diet (Skretting). Water temperature was maintained at 16°C. All experiments were reviewed and approved by the Institutional Animal Care and Use Committee at the University of New Mexico (protocol 19-200863-MC).

### 2.2. Sampling

Fish were sampled for histology at 340DD (mean weight 0.12g), 500DD (mean weight 0.21g) 700DD (mean weight 0.39g), 1000DD (mean weight 2.83g), 1400DD (mean weight 3.08g), and 1860DD (mean weight 3.34g). On the day of the sampling, fish were euthanized with an overdose of buffered MS-222 (250 mg/l, Syndel, catalog #Tricaine1G). Fish were weighed, and heads were removed behind the operculum. Heads were either fixed in 4% w/v PFA in PBS, or flash frozen in OCT (Sakura Finetek USA, catalog #4583).

### 2.3. Light Microscopy

Heads were fixed overnight in 4% w/v PFA in PBS at room temperature. Following decalcification in 10% w/v EDTA (pH 7.4) for 5 d, samples were moved to 70% v/v ethanol and processed in a tissue processor and embedded in paraffin in a sagittal orientation. For ages up to 1400DD, parasagittal paraffin sections (5 µm) were stained, while for 1865DD samples parasagittal cryosections (5 µm) were stained. Routine hematoxylin and eosin (H&E) stainings and Periodic Acid Schiff (PAS) staining (Millipore-Sigma, catalog #395B-1KT) were performed per manufacturer’s instructions. Large image acquisition was performed using a Leica Aperio AT2 slide scanner, and images were viewed using Leica ImageScope 12.4.6 software.

### 2.4. Immunofluorescence staining and confocal microscopy imaging

Prior to cryosectioning, fish heads sampled as described above, cut behind the operculum, were embedded in OCT and flash frozen. For all stainings, 5 µm-thick parasagittal cryosections were used. Sections were fixed for 3 minutes in 4% w/v PFA at room temperature, followed by blocking with Starting Block T20 blocking buffer (Thermo Fisher Scientific catalog #37539) for 13 minutes. All antibodies were diluted in PBST. Following antibody staining, all cell nuclei were labelled with DAPI (5µg/ml, Thermo Fisher Scientific, catalog #D1306). Samples were imaged using a Zeiss LSM 780 with Zen Blue software image stitching where applicable.

For immunostaining with rat anti-trout CD8α (Takizawa et al., 2011), cryosections were postfixed in ice-cold acetone, followed by 4% PFA, and labeled with non-affinity purified rat IgG anti-trout CD8α (1:80) overnight at 4°C, followed by AlexaFluor488-conjugated donkey anti-Rat IgG (1:300, Jackson Immunoresearch, Catalog #712-545-153) at room temperature for 2 h.

For immunostaining with mouse anti-trout MHC-II (Granja et al., 2015), cryosections were post-fixed in 4% PFA and labelled with mouse anti-trout MHC-II (2µg/ml) overnight at 4°C, followed by AlexaFluor488-conjugated donkey anti-mouse (Jackson Immunoresearch, Catalog #715-545-150) at room temperature for 2 h.

For triple staining to detect trout IgM^+^, IgT^+^ and CD4-2b^+^ cells, three 5 µm cryosections were post-fixed in 4% PFA, rinsed, blocked and labeled with mouse anti-trout IgM IgG1 antibody (1.4 µg/ml) (DeLuca et al., 1983), AlexaFluor555-labeled mouse anti-trout IgT (Zhang et al., 2010) and guinea pig anti-Trout CD4-2b overnight at 4°C, followed by AlexaFluor488 goat anti-guinea pig IgG (1:300, Jackson Immunoresearch, Catalog #106-545-003) and Cy5-labelled goat anti-mouse IgG1 (1:300, Jackson Immunoresearch, catalog no. 115-175-166) at room temperature for 2 hours.

Images of the O-NALT (N=3-5) were acquired with a Zeiss LSM780 confocal microscope. Images were viewed and analyzed using Fiji. Approximate lymphocyte composition at 1400DD was determined by measuring the area of the O-NALT in Fiji, then counting the number of positive cells within that area to determine cells/μm^2^, which was then used to determine the approximate proportions. Positive cells were defined as a clear DAPI nucleus surrounded by positive membrane signal.

### 2.5. Gene expression analysis

Tissues were homogenized in TRIZol (Invitrogen, catalog #15596026) using tungsten carbide beads (3 mm, Qiagen catalog #69997) and shaking (300 times per min) in a Tissuelyser per manufacturer’s instructions for RNA extraction. The RNA pellet was washed in 80% v/v ethanol, air-dried and resuspended in RNase-free water. RNA quality and quantity were determined by NanoDrop (Thermo Fisher Scientific). For all genes, expression levels were determined using the Luna Universal One-Step RTqPCR kit (New England Biolabs catalog #E3005X) per manufacturer’s instructions. For each reaction, 500 ng of template RNA was used. Primer sequences and annealing temperatures used are listed in Table 1. Reactions were performed using the detection system BioRad CFX96 C1000 Touch real-time PCR (Bio-Rad).

**Table 1:**
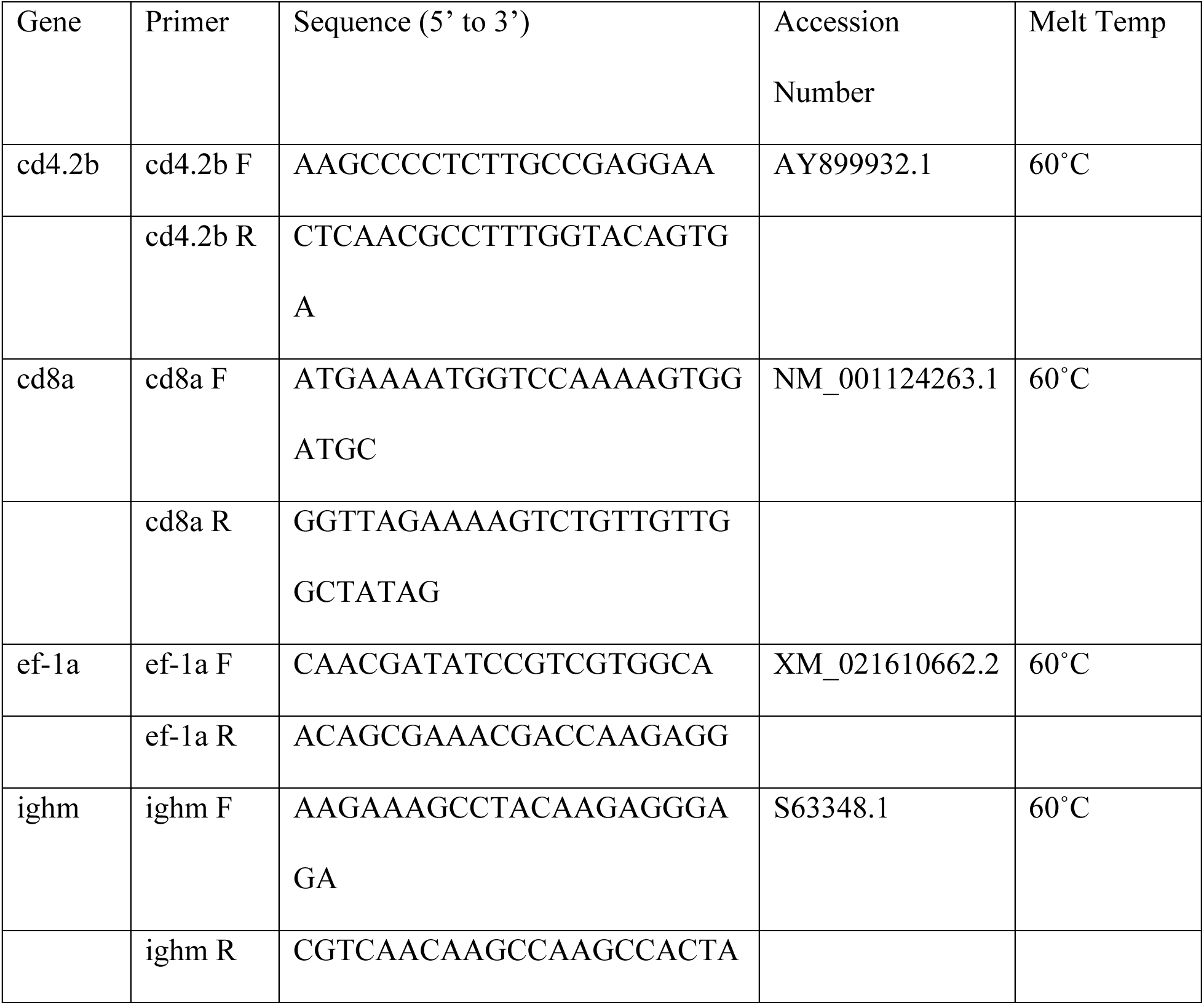
qPCR Primers.

### 2.6. Statistical analyses

For gene expression data, results were analyzed by the Pfaffl method (Pfaffl, 2001) and expression was relative to *ef1a* as previously performed (Garcia et al., 2022). Statistical differences between groups were done by one-way ANOVA, with a P<0.05 considered significant. All data were analyzed in Prism v.9.

## 3. Results

### 3.1. Histological changes in the nasal cavity of rainbow trout during development

Histological observations of H&E-stained paraffin sections indicated that the full O-NALT structure with the characteristic bulging appearance of lymphocytes pushing into the basal membrane (Garcia et al., 2022) is present by 1400DD. At 340DD, few to no intraepithelial lymphocytes were apparent in the lining of the nasal cavity (Fig. 1A), which is only composed of epithelial cells. By 500DD, reticular epithelial cells were present between the basal lamina and the most superficial epithelial cells, but there were still no lymphocytes embedded in them as observed in developed O-NALT (Fig. 1C). By 700DD (Fig. 1E) a few scattered lymphocytes were observed intraepithelially in the nasal cavity, but they did not appear to be clustered yet. At 1000DD (Fig 1G), the bulging structure of the O-NALT had not yet formed, and there were not yet clusters of lymphocytes present. However, there are numerous mucus-producing cells scattered throughout the lining of the nasal cavity and visible even by H&E staining. By 1400 DD (Fig. 1I), clusters of intraepithelial lymphocytes were scattered throughout the nasal cavity embedded in reticulated epithelial cells, with particularly large clusters present opposite the tips of the lamellae, as we previously observed in immunologically mature fish (Garcia et al., 2022). By 1860DD, clusters of lymphocytes were present at the bases of the lamellae, as well as directly opposite the lamellae and the O-NALT structure was larger than observed in 1400DD fish (Fig 1K).

**Figure 1:**
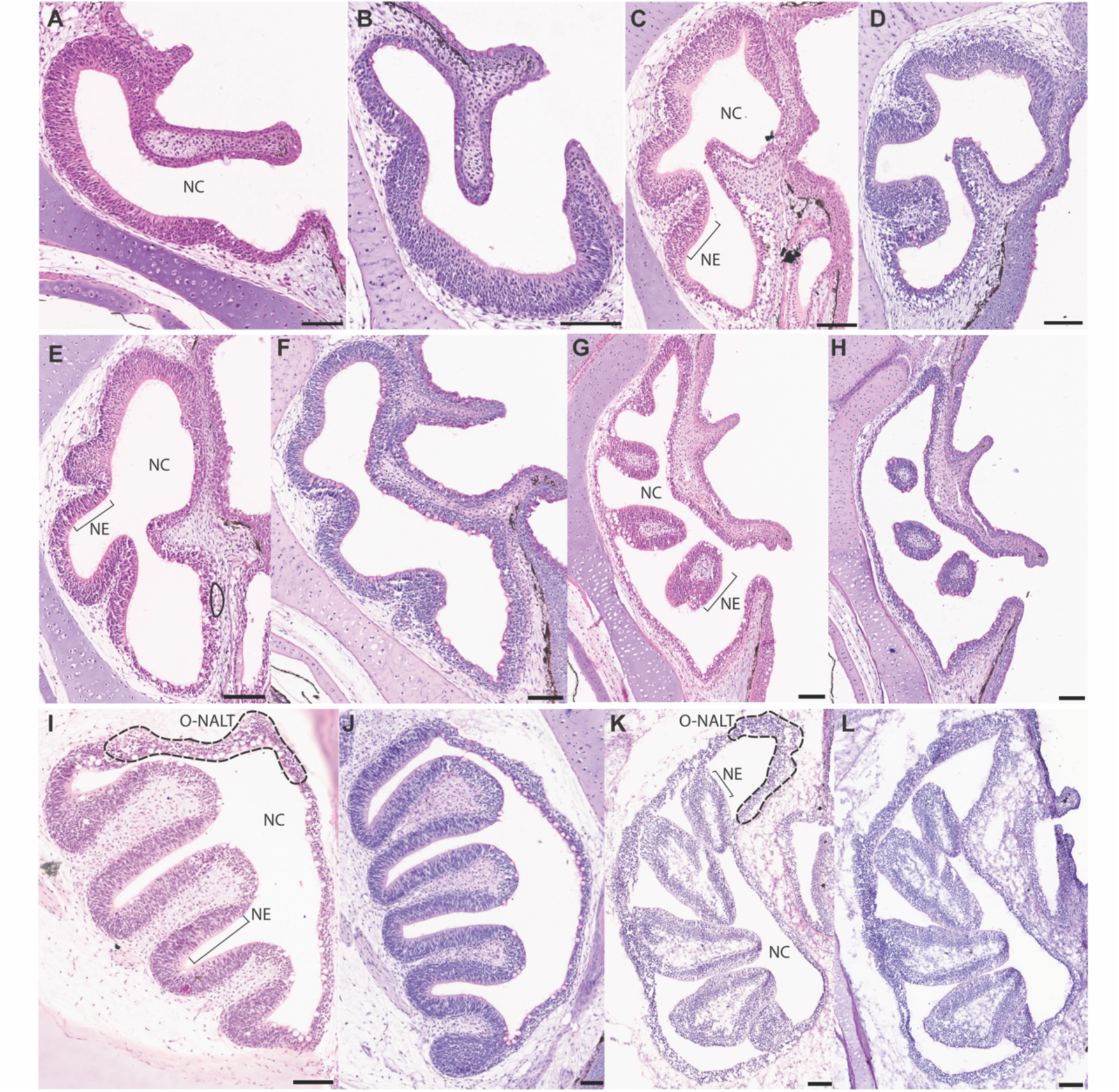
Histomormorphological changes that lead to O-NALT development in rainbow trout. Representative H&E stainings at 340DD (A), 500DD (C), 700DD (E), and 1000DD (G) do not show any obvious bulging lymphocyte rich structure indicative of the O-NALT. By 1400DD (I), this structure is visible and continues to be present at 1860DD (K). PAS staining at 340DD (B) shows very few goblet cells in the nasal cavity, with increased numbers at 500DD (D), and continuing to 700DD (F), and 1000DD (H). By 1400DD, numerous goblet cells are visible superficial to the lymphocytes present in the O-NALT, at a far higher density than in other areas of the nasal cavity lining. At 1860DD, PAS^+^ cells are no longer present in the nasal cavity. All scale bars are 100μm.

PAS staining showed very few PAS^+^ cells present inside the nasal cavity at 340DD, although some were present on the skin (Fig. 1B). PAS^+^ cells were present in the nasal cavity by 500DD (Fig. 1D). There was a noticeable increase in PAS^+^ cell numbers between 700DD (Fig. 1F) and 1000DD (Fig. 1H). At 1400DD, the characteristic O-NALT bulge showed an increased density of goblet cells covering the O-NALT. The O-NALT, at this developmental stage, measures ∼30µm deep by ∼150µm wide at its largest possible area. At 1860DD, we did not observe PAS^+^ cells in the trout nasal cavity. Initially, we thought this may be an issue with the staining or an artifact. However, clear PAS^+^ cells were present nearby in the oral cavity on the same slide, and what appeared morphologically by H&E to be goblet cells were indeed present in the nasal cavity. As such, we concluded that the lack of PAS^+^ cells in the nasal cavity at 1800DD is most likely due to differences in mucin composition, an observation that warrants further investigation. In the olfactory epithelium, there were few goblet cells present by PAS staining at all developmental time points, and as previously reported, most goblet cells present on the lamellae in the nasal cavity are present at either the bases or the tips where the mucosal epithelium and not neuroepithelium is located (Sepahi et al., 2016).

### 3.2. Ontogenic changes of B and T cells in O-NALT

Immunofluorescence staining indicated that the first CD4-2b^+^ and IgM^+^ cells do not appear in the nasal cavity until 1000DD (Fig. 2A-D). At 1000DD, both CD4-2b^+^ and IgM^+^ cells were observed around the entrance to the nasal cavity, in the same areas where they have previously been reported (Garcia et al., 2022). While we observed both CD4-2b^+^ and IgM^+^ cells in the nasal cavity at 1000DD, the observed clusters of IgM^+^ cells forming the characteristic bulge of the O-NALT were not present until 1400DD. At 1400DD, we observed large numbers of CD4-2b^+^ cells present intraepithelially, generally close to the basement membrane. In contrast, clusters of IgM^+^ cells were also present, but they tended to be more superficially located compared to CD4-2b^+^ cells. By 1800DD, CD4-2b^+^ and IgM^+^ cells were both present at high densities throughout the O-NALT. While CD4-2b^+^ cells were relatively evenly distributed throughout the O-NALT, IgM^+^ cells were present in distinct clusters (Fig 2F, I). In the olfactory lamellae, CD4-2b^+^ and IgM^+^ cells were not found in the olfactory epithelium itself, but rather are most often seen in the lamina propria of the lamellae (Fig. 2D-F).

**Figure 2:**
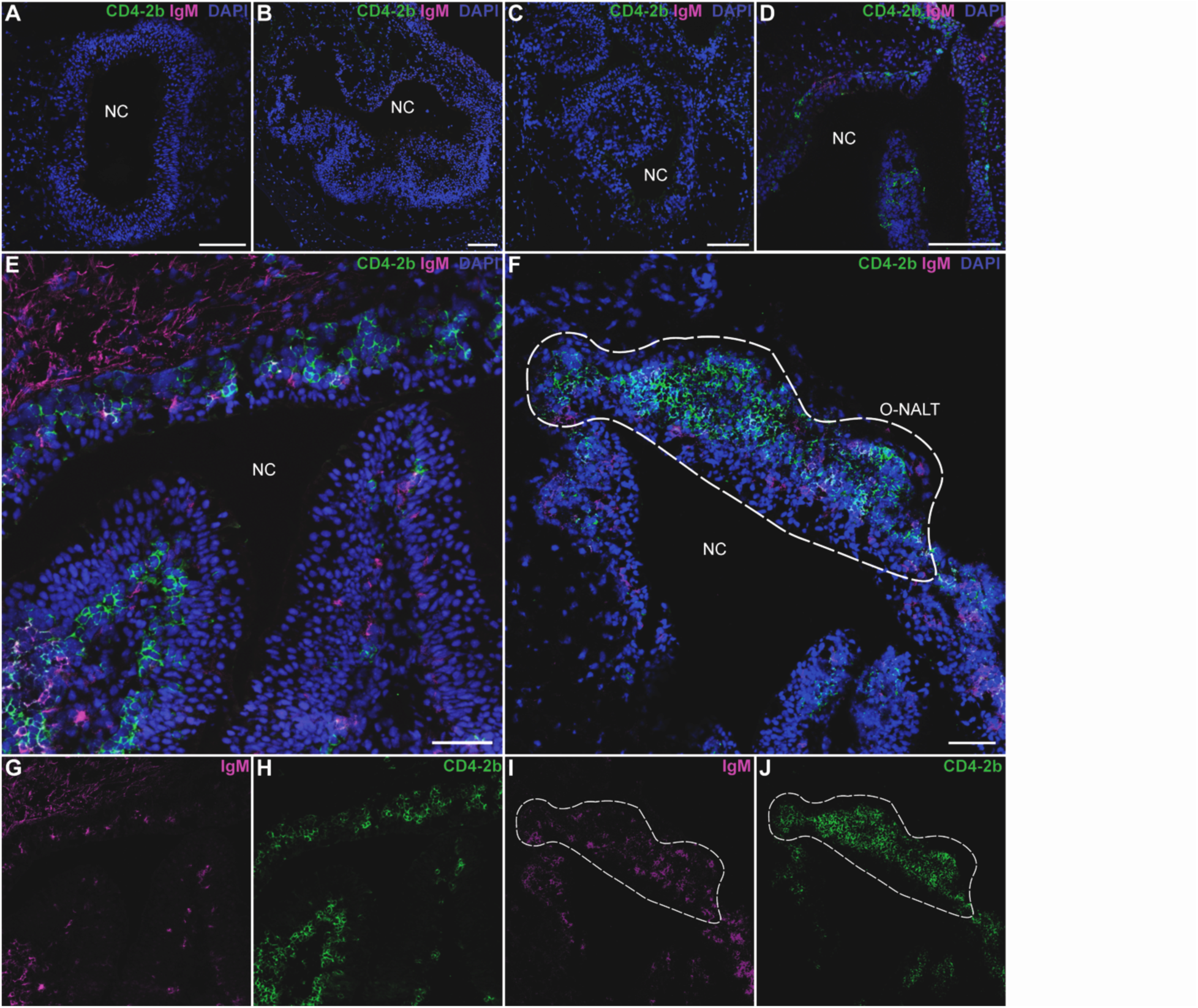
Ontogeny of IgM^+^ and CD4-2b^+^ cells are the trout nasal cavity. Confocal images of sagittal cryosections of rainbow trout nasal cavity stained with anti-CD4-2b (green), anti-IgM (magenta) and DAPI (blue) at 340DD (A), 500DD (B), 700DD (C), 1000DD (D), 1400D (E), and 1860DD (F). Single channel images of 1400DD anti-IgM (G) and anti-CD4-2b (H) stains show some clusters of CD4-2b^+^ and IgM^+^ cells. Single channel images of 1860DD anti-IgM (I) and anti-CD4-2b (J) stains show large clusters of CD4-2b^+^ and IgM^+^ cells indicative of the O-NALT. Scale bars 50µm (D, E, G, and H), and 100µm A, B, C, and F). NC, Nasal cavity; dotted lines in F, I, J indicate the O-NALT.

CD8α staining indicated a few, scattered CD8α^+^ cells present in the nasal cavity by 700DD, prior to the presence of either CD4-2b^+^ T cells or IgM^+^ B cells, and by 1400DD, clusters of CD8α^+^ cells were observed at the intraepithelial level. By 1860DD, more CD8α^+^ cells were present in the bulging structure characteristic of the O-NALT, but they were not as abundant as CD4-2b^+^ cells (Fig. 3F). At all time points, few CD8α^+^ T cells were present in the olfactory epithelium, and most CD8α^+^ cells present in the lamellae of the nasal cavity are in the lamina propria or on the tips of the lamellae (Fig. 3E,F).

**Figure 3:**
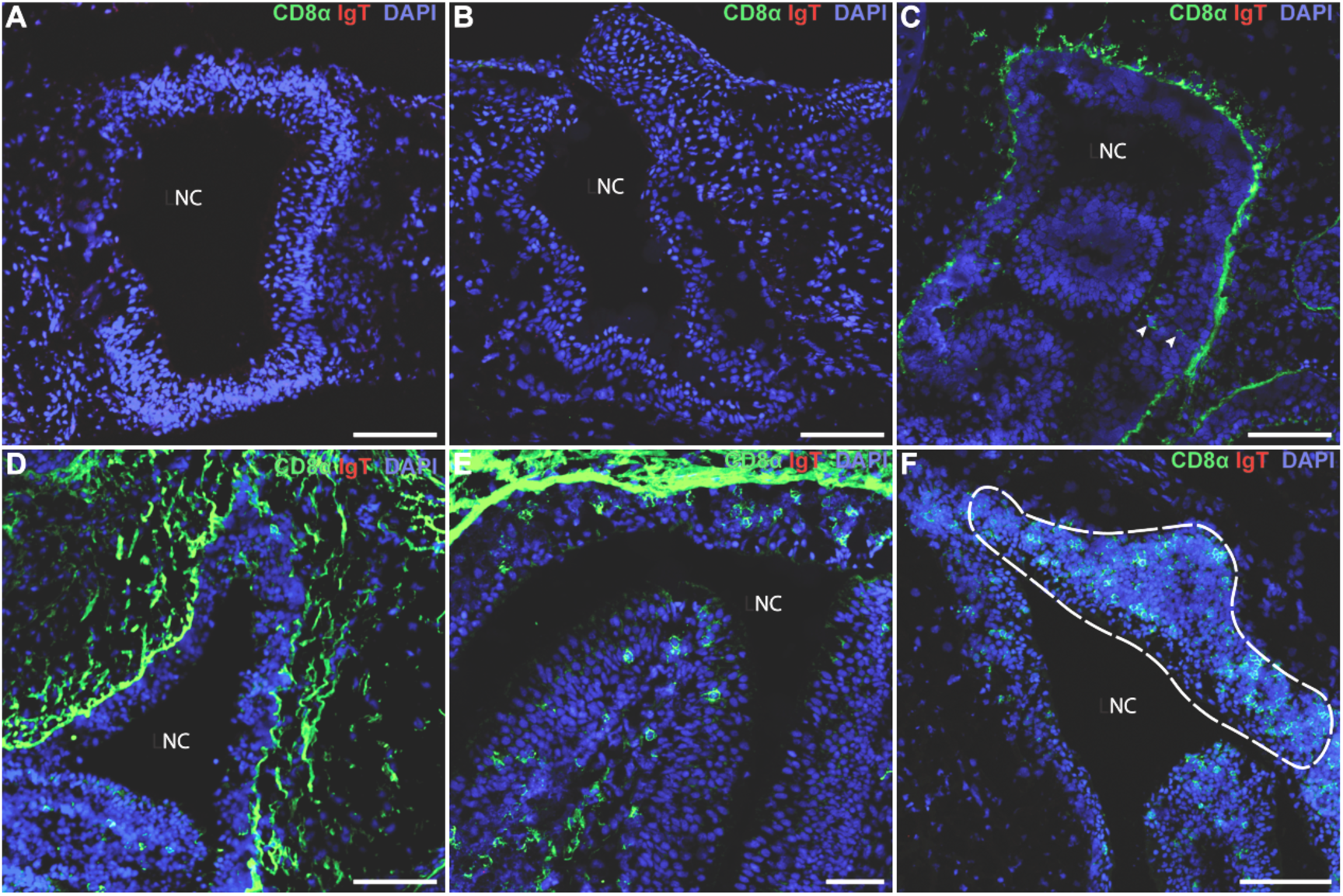
Ontogeny of CD8α^+^ and IgT^+^ cells in the trout nasal cavity. At 340DD (A) and 500DD (B), CD8α^+^ cells (green) are not yet present in the nasal cavity. By 700DD (C), CD8α^+^ cells can be observed intraepithelially (indicated by white arrows), but they are not yet present in clusters. By 1000DD (D), more CD8α^+^ cells are present intraepithelially, but clusters are not apparent at an intraepithelial level until 1400DD (E). By 1860DD (F), clusters of CD8α^+^ cells can be observed in the characteristic bulging O-NALT structure. Scale bars are 100μm (A-D), 50μm (E), and 200μm (F). NC, Nasal Cavity. Dotted line in F indicates classical O-NALT structure.

Interestingly, we observed no IgT^+^ B cells in the nasal cavity at any time point in this study. While some IgT^+^ B cells were present on the skin, IgT^+^ B cells were not observed anywhere in the nasal cavity, including the O-NALT, olfactory epithelium, and lymphoepithelial lining, even by 1860DD (Fig. 3F).

By 1400DD, the O-NALT is composed of approximately 20% IgM^+^ B cells, 67% CD4-2b^+^ T cells, and 13% CD8α^+^ T cells, compared to 24% IgM^+^ B cells, 56%CD4-2b^+^ T cells, 16% CD8α^+^ T cells, and 4% IgT^+^ B cells at the 30g stage (Garcia et al., 2022).

### 3.3. Ontogeny of MHC-II^+^ cells in the rainbow trout nasal cavity

MHC-II staining of the O-NALT showed low-level expression in epithelial cells as early as 500 DD (Fig. 4B), in line with what has been previously reported (Sepahi et al., 2016). High level expression of MHC-II, putatively in professional antigen presenting cells based on their size and shape (Fig. 4D, F), was first observed at significant levels at 1000DD. By 1400DD high density of MHC-II^low^ and MHC-II^high^ expressing cells was observed scattered throughout the O-NALT structure. These included both round, lymphocyte-like cells as well as larger cells with dendritic morphology which could correspond to putative macrophages and/or dendritic cells. Additionally, the covering epithelium of the O-NALT structure also contained MHC-II^+^ cells including some oval, large, MHC-II^high^ cells which may be goblet cells (Fig. 4D, F). The presence of this bulging structure continued at 1860DD (Fig. 4G), with an increased density of MHC-II^+^ cells relative to 1400DD.

**Figure 4:**
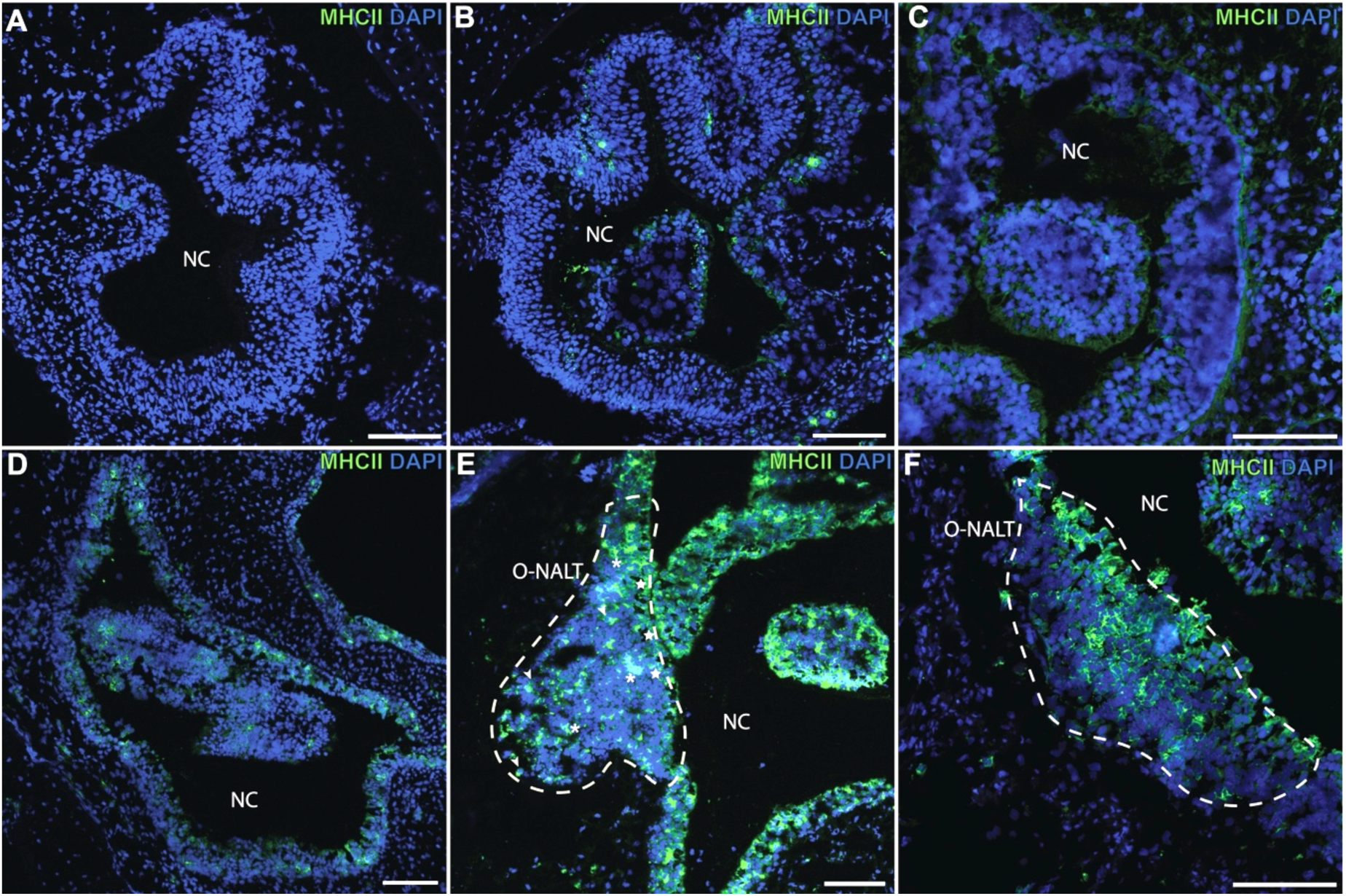
Ontogenic changes in MHC-II^+^ cells in the trout nasal cavity. Confocal microscopy images of developing trout nasal cavity cryosections at 340DD (A) where there is no MHC-II^+^ cells present in the nasal cavity. 500DD (B) where some scattered MHC-II^+^ cells are present, with low level expression in epithelial cells. 715DD (C) low level MHC-II expression is seen across almost all epithelial cells in the nasal cavity. 1000DD (D) large numbers of MHC-II^+high^ cells are observed throughout the nasal cavity and low level MHC-II expression is still visble in epithelial cells. 1400DD (E) MHC-II^+high^ cells with different morphology and MHC-II^+low^ cells are observed in the O-NALT bulge. At 1860DD the number of MHC-II^+^ cells in the O-NALT continues to increase and the characteristic bulging structure is larger. Scale bars are 100µm. NC, Nasal cavity. Dotted lines in E and F indicate the O-NALT. In E, Asterisks denote MHC-II^high^ putative antigen presenting cells. Arrowheads denote MHC-II^+^ putative lymphocytes. Stars denote MHC-II^+^ oval shaped cells, putative goblet cells.

### 3.4 Whole-body changes in immune gene expression

qPCR analyses indicated that whole body expression levels of *cd8a* and *cd4-2b* followed patterns similar to those observed by immunofluorescent staining in the nasal cavity (Fig. 5). Relative expression of *cd4-2b* at 750DD was 16-fold higher than that of the first developmental time point studied (Fig. 5A). By 860DD, *cd4-2b* relative expression decreased slightly (13.7-fold increase relative to 300DD).

**Figure 5:**
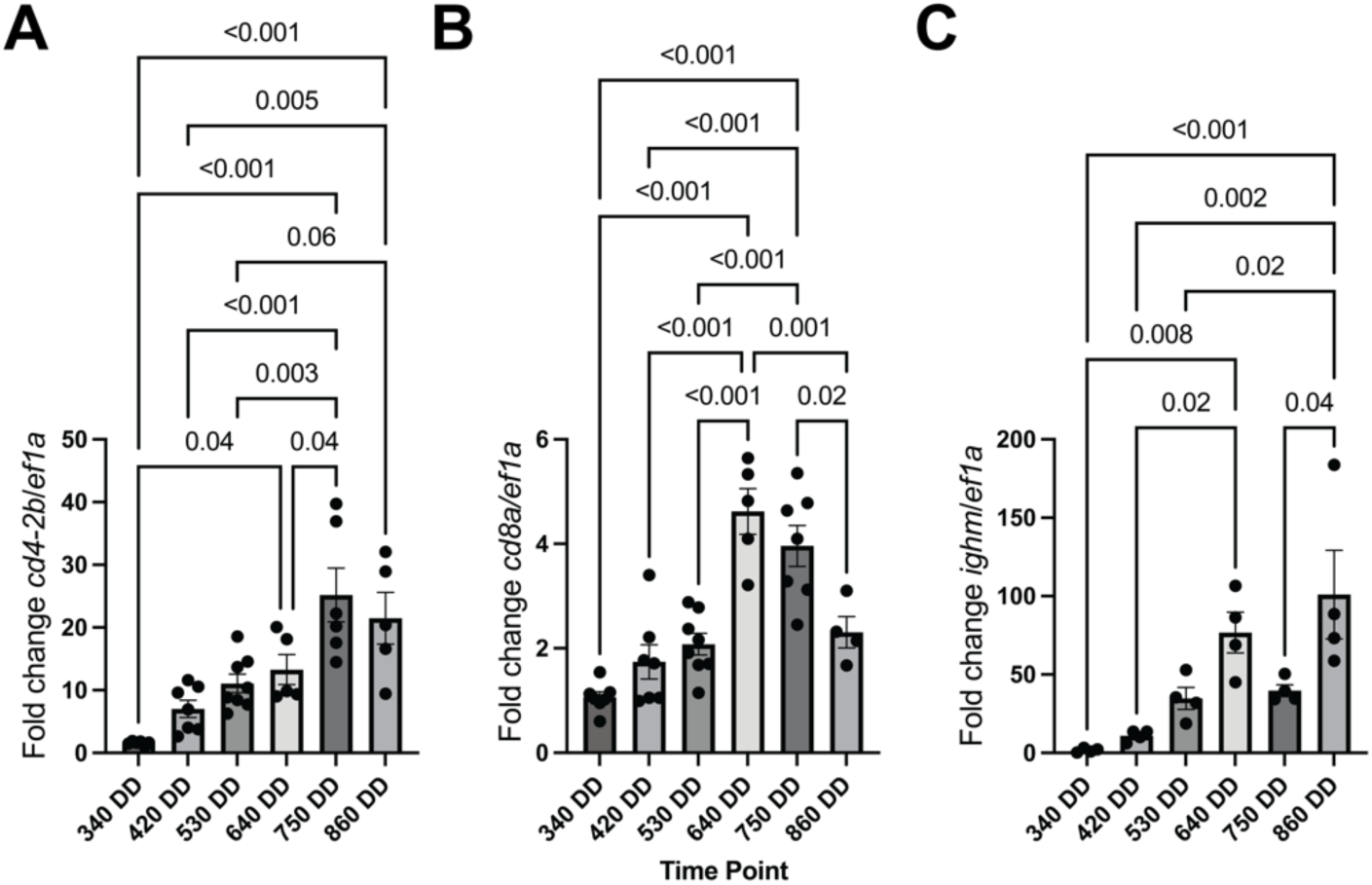
Whole body immune gene expression changes during trout ontogeny. Relative expression of *cd4-2b* to *ef1a* (A), *cd8a* to *ef1a* (B), and *ighm* to *ef1a* (C) calculated using the Pfaffl method. One-way ANOVA was used to determine differences between time points.

**Figure 6:**
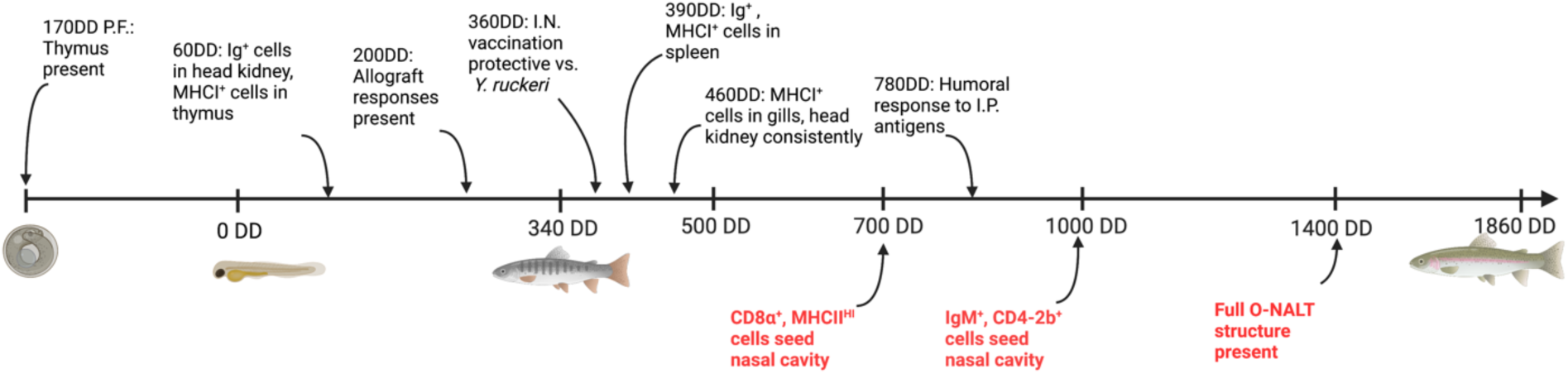
Timeline of the developmental trajectory of adaptive immunity in *O. mykiss.* Schematic summary of the findings from this study as well as previous published studies. Developmental timepoint pre-hatching is indicated in days pre-hatch at 14°C. Developmental time points after hatching are indicated in degree days. Indications below the timeline in red text are conclusions based on results in this paper. Indications above the timeline in black text are taken from the following: (Fischer et al., 2005; Razquin et al., 1990; Salinas et al., 2015; Tatner, 1986; Tatner and Manning, 1983). DD, Degree Days; I.N., Intranasal; I.P., Intraperitoneal; PF, Post-fertilization. Unless otherwise noted, degree days are measured from the time of hatching. Figure was created in Biorender.

In contrast, relative expression of *cd8a* at 640DD (Fig. 5B) was 4.3-fold higher than that of 300DD fish. After 640DD, *cd8a* relative expression decreased precipitously: by 860DD, *cd8a* relative expression decreased to only 2.2-fold higher than at 340DD.

Expression of *ighm* steadily increased from 340DD to 640DD, when its relative expression was 76.7-fold higher than in 340DD fish (Fig. 5C). At 750DD, *ighm* relative expression was 40-fold that of 340DD fish. By 860DD, *ighm* relative expression peaked to 101-fold that of 340DD fish.

## 4. Discussion

An emerging body of work indicates that the architecture of the mucosal immune system in teleosts is much more complex than originally thought with several organized or semi-organized MALT found in several species (Dalum et al., 2021; Garcia et al., 2022; Haugarvoll et al., 2008; Koppang et al., 2010; Resseguier et al., 2023). Yet, despite the importance of the O-MALT microenvironment in the initiation of adaptive immune responses at mucosal tissues, when and how these structures form during teleost ontogeny is largely unknown. The present study focused on one O-MALT structure, the O-NALT, and mapped the cellular changes that occur in this structure from 340DD to 1800DD in rainbow trout.

NALT development in rodent models is composed of a series of cellular seeding events, during which different cell types dominate and are then replaced. In rats, CD8^+^ T cells are abundant in NALT early in development, with CD4^+^ T cells becoming predominant around 45 days old (Sosa and Roux, 2004). Despite this early predominance of CD8^+^ T cells, by 14 days, distinct B and T cell zones are present in the rat NALT (van Poppel et al., 1993). This distinct cellular cascade, with CD8^+^ T cells leading, potentially playing an effector role, followed by CD4^+^ T cells and B cells, potentially playing an inductor role, suggests that changes to NALT cellular composition may be related to functional changes. Similarly, the present study reveals a cascade of events that leads to a fully formed O-NALT structure by 1400DD. Specifically, in trout, CD8α^+^ cells appear in the O-NALT well before B cells, around 500DD, aligning with evidence from rats (Sosa and Roux, 2004). Furthermore, our finding that CD8α^+^ T cells seed the nasal cavity prior to B cells aligns with findings from the GALT of common carp (*Cyprinus carpio*) (Sosa and Roux, 2004). In support, previous studies found expression of *cd8a* by RT-PCR in trout as early as 84DD post-fertilization, about 2 weeks prior to hatching, highlighting the importance of this molecule in early life immune defense (Fischer et al., 2005). The early CD8α ^+^ T cells detected in the nasal cavity in the present study may represent innate CD8α ^+^ T cells, as we have previously reported that they are critical in very rapid, trout nasal immune responses to viruses (Sepahi et al., 2019).

Interestingly, our previous work (Garcia et al., 2022) as well as the present study suggest that IgT^+^ B cells contribute minimally to the composition of O-NALT and suggest that IgT found in nasal mucus of immunologically mature fish (Tacchi et al., 2014) is likely coming from the scattered IgT^+^ B cells found in the olfactory epithelium. However, further studies are warranted, especially in immunized animals, to verify whether IgT^+^ B cells participate in the immune response within the O-NALT structure.

In mammalian O-MALT structures, specialized antigen sampling cells expressing MHC-II are found in the modified epithelium that covers O-MALT such as Peyer’s patches, tonsils, and NALT (Date et al., 2017; Kiyono and Fukuyama, 2004; Wosen et al., 2018). In teleosts, MHC-II expression has been examined in the ILT of Atlantic salmon (Austbo et al., 2014), and the ALT and NEMO of zebrafish (Dalum et al., 2021; Resseguier et al., 2023). In the ILT, MHC-II-expressing cells are found scattered within the structure. Using several transgenic lines, the different MHC-II^+^ cells in the zebrafish ALT and NEMO were resolved. These include MHC-II^low^ reticular epithelial cells and MHC-II^high^ professional antigen presenting cells including dendritic cells, macrophages, lymphocytes and granulocytes (Dallum et al., 2021; Resseguier et al., 2023). Although we did not phenotype the different subsets of MHC-II^+^ cells in the trout O-NALT, based on morphology, we observed low expressing cells with epithelial morphology as well as high expressing cells with reticular morphology which are putative dendritic cells and macrophages as well as small round cells, which are likely B cells. Additionally, the covering epithelium contained some large oval positive cells with high levels of expression that need further characterization. Interestingly, while epithelial MHC-II^low^ cells were found early during ontogeny, putative professional antigen presenting cells were only observed from 1000DD onwards, potentially indicating functional differences in the ability of antigens to induce specific immune responses in the nasal cavity of young rainbow trout.

Microbial antigens continuously stimulate MALT, yet, their development in mammals is not dependent on antigenic stimulation but developmentally programed (Drayton et al., 2006). Tertiary lymphoid organs (TLOs), on the other hand, are formed upon antigenic exposure in different body sites (Drayton et al., 2006). In fish, antigen stimulation at mucosal surfaces at the time of first feeding coincides with the expression of a number of adaptive immune markers like MHC-I (Fischer et al., 2005). The timeline described and methods used allow us to ascertain whether O-NALT development in teleost fish is developmentally programmed or whether microbial stimulation plays a role in accelerating development as seen in mammalian models (Krege et al., 2009). This question may be best answered in zebrafish or rainbow trout germ-free models (Perez-Pascual et al., 2021; Rawls et al., 2004), an exciting future direction to this study. Investigation of the ontogeny of other lymphoid structures in trout such as ILT, ALT and NEMO will elucidate if they also develop along the same trajectory, which would help determine whether similar developmental triggers or microbial stimulation are responsible for the formation of all these MALTs.

In mammalian models, NALT lymphoid structures develop by different mechanisms than other organized MALT (Fukuyama et al., 2002). Most lymphoid tissues, including other mucosal tissues like Peyer’s patches, develop in a lymphotoxin-α (LTα), RORγ dependent manner (Randall et al., 2008). However, NALT successfully develops in both LTα^−/−^ and RORγ^−/−^ mice, albeit with a smaller, disorganized structure (Harmsen et al., 2002). While independent of LTα and RORγ, NALT development is dependent on ID2, a transcription factor inhibition protein, known to have numerous roles in lymphoid tissue development (Fukuyama et al., 2002). ID2 drives differentiation of a class of CD3^−^CD4^+^CD45^+^ cells which seed the NALT early during development in mice. These CD3^−^CD4^+^CD45^+^ cells are both necessary and sufficient for NALT development in mouse models (Fukuyama et al., 2002). While teleost fish lack LTα, they possess functional and highly conserved ID2, which is believed to be involved in the formation of lymphoid organs like the thymus (Lane et al., 2012). In African lungfish, NALT lymphoid aggregates also express ID2 (Tacchi et al., 2015). The high conservation of ID2, and its known role in NALT development in mice, indicates that patterns of NALT development may be conserved across a broad range of taxa.

The limitations of the present study surround two main areas: first, we do not have direct evidence that morphological, cellular and gene expression changes at 1400DD correlate with functional adaptive immune responses since we did not perform any immunization studies (Garcia et al., 2022). Second, we did not evaluate B cell repertoire changes in O-NALT in control and immunized animals, but ongoing work in our laboratory is focusing on this question.

Beyond increasing fundamental understanding of mucosal immunity in teleosts, our findings have important implications for the rationale design of mucosal vaccines in farmed fish. Our previous work has demonstrated that intranasal vaccination induces high levels of protection as early as 360DD in rainbow trout, protection measured by challenge 28 days post-vaccination (Salinas et al., 2015). Thus, it is clear that nasal vaccines are effective even when O-NALT is not fully formed yet, potentially mediated by protection conferred by diffuse NALT. However, it is possible that delaying the age of primary vaccination to a timepoint where O-NALT is formed may lead to longer-lasting, higher affinity specific immune responses, a question we are currently exploring in our laboratory. Delaying the age of vaccination, however, may leave animals unprotected when they are the most susceptible to some pathogens, complicating the decision as to when it is optimal to vaccinate.

## 5. Conclusions

The present study provides the first detailed characterization of the ontogeny of a teleost O-MALT, the O-NALT. We identify a unique sequence of events leading to the full formation of O-NALT at 1400DD. CD8α^+^ T cells, CD4-2b^+^ T cells and IgM^+^ B cells seed O-NALT in that order overall recapitulating quite well whole-body changes in gene expression. Future studies will determine whether timing of nasal vaccination to O-NALT emergence brings any substantial benefits in terms of the quality and duration of the specific immune response.

## Acknowledgements

This work was funded by the US Department of Agriculture USDA NIFA award # 2019-05906 to IS. Authors thank Drs. Sunyer, Takizawa and Tafalla for providing the primary antibodies used in this study. The authors thank the UNM ROSE Scholar program for the support provided to C. Martinez.

## References

Aas, I.B., Austbo, L., Falk, K., Hordvik, I., Koppang, E.O., 2017. The interbranchial lymphoid tissue likely contributes to immune tolerance and defense in the gills of Atlantic salmon. Dev Comp Immunol 76, 247–254.

Austbo, L., Aas, I.B., Konig, M., Weli, S.C., Syed, M., Falk, K., Koppang, E.O., 2014. Transcriptional response of immune genes in gills and the interbranchial lymphoid tissue of Atlantic salmon challenged with infectious salmon anaemia virus. Dev Comp Immunol 45, 107–114.

Buchmann, K., 2022. The Ontogeny of the Fish Immune System, in: Buchmann, K., Secombes, C.J. (Eds.), Principles of Fish Immunology : From Cells and Molecules to Host Protection. Springer International Publishing, Cham, pp. 495–510.

Dalum, A.S., Austbo, L., Bjorgen, H., Skjodt, K., Hordvik, I., Hansen, T., Fjelldal, P.G., Press, C.M., Griffiths, D.J., Koppang, E.O., 2015. The interbranchial lymphoid tissue of Atlantic Salmon (Salmo salar L) extends as a diffuse mucosal lymphoid tissue throughout the trailing edge of the gill filament. J Morphol 276, 1075–1088.

Dalum, A.S., Griffiths, D.J., Valen, E.C., Amthor, K.S., Austbo, L., Koppang, E.O., Press, C.M., Kvellestad, A., 2016. Morphological and functional development of the interbranchial lymphoid tissue (ILT) in Atlantic salmon (Salmo salar L). Fish Shellfish Immunol 58, 153–164.

Dalum, A.S., Kraus, A., Khan, S., Davydova, E., Rigaudeau, D., Bjorgen, H., Lopez-Porras, A., Griffiths, G., Wiegertjes, G.F., Koppang, E.O., Salinas, I., Boudinot, P., Resseguier, J., 2021. High-Resolution, 3D Imaging of the Zebrafish Gill-Associated Lymphoid Tissue (GIALT) Reveals a Novel Lymphoid Structure, the Amphibranchial Lymphoid Tissue. Front Immunol 12, 769901.

Date, Y., Ebisawa, M., Fukuda, S., Shima, H., Obata, Y., Takahashi, D., Kato, T., Hanazato, M., Nakato, G., Williams, I.R., Hase, K., Ohno, H., 2017. NALT M cells are important for immune induction for the common mucosal immune system. Int Immunol 29, 471–478.

DeLuca, D., Wilson, M., Warr, G.W., 1983. Lymphocyte heterogeneity in the trout, Salmo gairdneri, defined with monoclonal antibodies to IgM. Eur J Immunol 13, 546–551.

Drayton, D.L., Liao, S., Mounzer, R.H., Ruddle, N.H., 2006. Lymphoid organ development: from ontogeny to neogenesis. Nat Immunol 7, 344–353.

Fischer, U., Dijkstra, J.M., Kollner, B., Kiryu, I., Koppang, E.O., Hordvik, I., Sawamoto, Y., Ototake, M., 2005. The ontogeny of MHC class I expression in rainbow trout (Oncorhynchus mykiss). Fish Shellfish Immunol 18, 49–60.

Fukuyama, S., Hiroi, T., Yokota, Y., Rennert, P.D., Yanagita, M., Kinoshita, N., Terawaki, S., Shikina, T., Yamamoto, M., Kurono, Y., Kiyono, H., 2002. Initiation of NALT organogenesis is independent of the IL-7R, LTbetaR, and NIK signaling pathways but requires the Id2 gene and CD3(−)CD4(+)CD45(+) cells. Immunity 17, 31–40.

Garcia, B., Dong, F., Casadei, E., Resseguier, J., Ma, J., Cain, K.D., Castrillo, P.A., Xu, Z., Salinas, I., 2022. A Novel Organized Nasopharynx-Associated Lymphoid Tissue in Teleosts That Expresses Molecular Markers Characteristic of Mammalian Germinal Centers. J Immunol 209, 2215–2226.

Granja, A.G., Leal, E., Pignatelli, J., Castro, R., Abos, B., Kato, G., Fischer, U., Tafalla, C., 2015. Identification of Teleost Skin CD8alpha+ Dendritic-like Cells, Representing a Potential Common Ancestor for Mammalian Cross-Presenting Dendritic Cells. J Immunol 195, 1825–1837.

Grontvedt, R.N., Espelid, S., 2003. Immunoglobulin producing cells in the spotted wolffish (Anarhichas minor Olafsen): localization in adults and during juvenile development. Dev Comp Immunol 27, 569–578.

Harmsen, A., Kusser, K., Hartson, L., Tighe, M., Sunshine, M.J., Sedgwick, J.D., Choi, Y., Littman, D.R., Randall, T.D., 2002. Cutting edge: organogenesis of nasal-associated lymphoid tissue (NALT) occurs independently of lymphotoxin-alpha (LT alpha) and retinoic acid receptor-related orphan receptor-gamma, but the organization of NALT is LT alpha dependent. J Immunol 168, 986–990.

Haugarvoll, E., Bjerkas, I., Nowak, B.F., Hordvik, I., Koppang, E.O., 2008. Identification and characterization of a novel intraepithelial lymphoid tissue in the gills of Atlantic salmon. J Anat 213, 202–209.

Kiyono, H., Fukuyama, S., 2004. NALT-versus Peyer’s-patch-mediated mucosal immunity. Nat Rev Immunol 4, 699–710.

Koppang, E.O., Fischer, U., Moore, L., Tranulis, M.A., Dijkstra, J.M., Kollner, B., Aune, L., Jirillo, E., Hordvik, I., 2010. Salmonid T cells assemble in the thymus, spleen and in novel interbranchial lymphoid tissue. J Anat 217, 728–739.

Krege, J., Seth, S., Hardtke, S., Davalos-Misslitz, A.C., Forster, R., 2009. Antigen-dependent rescue of nose-associated lymphoid tissue (NALT) development independent of LTbetaR and CXCR5 signaling. Eur J Immunol 39, 2765–2778.

Lane, P.J., Gaspal, F.M., McConnell, F.M., Withers, D.R., Anderson, G., 2012. Lymphoid tissue inducer cells: pivotal cells in the evolution of CD4 immunity and tolerance? Front Immunol 3, 24.

Loken, O.M., Bjorgen, H., Hordvik, I., Koppang, E.O., 2020. A teleost structural analogue to the avian bursa of Fabricius. J Anat 236, 798–808.

Miller, R.D., Salinas, I., 2020. Phylogeny of the mucosal immune system, Principles of Mucosal Immunology. Garland Science, pp. 23–32.

Perez-Pascual, D., Vendrell-Fernandez, S., Audrain, B., Bernal-Bayard, J., Patino-Navarrete, R., Petit, V., Rigaudeau, D., Ghigo, J.M., 2021. Gnotobiotic rainbow trout (Oncorhynchus mykiss) model reveals endogenous bacteria that protect against Flavobacterium columnare infection. PLoS Pathog 17, e1009302.

Pfaffl, M.W., 2001. A new mathematical model for relative quantification in real-time RT-PCR. Nucleic Acids Res 29, e45.

Picchietti, S., Terribili, F.R., Mastrolia, L., Scapigliati, G., Abelli, L., 1997. Expression of lymphocyte antigenic determinants in developing gut-associated lymphoid tissue of the sea bass Dicentrarchus labrax (L.). Anat Embryol (Berl) 196, 457–463.

Randall, T.D., Carragher, D.M., Rangel-Moreno, J., 2008. Development of secondary lymphoid organs. Annu Rev Immunol 26, 627–650.

Rawls, J.F., Samuel, B.S., Gordon, J.I., 2004. Gnotobiotic zebrafish reveal evolutionarily conserved responses to the gut microbiota. Proc Natl Acad Sci U S A 101, 4596–4601.

Razquin, B.E., Castillo, A., Lopez-Fierro, P., Alvarez, F., Zapata, A., Villena, A.J., 1990. Ontogeny of IgM-producing cells in the lymphoid organs of rainbow trout, Salmo gairdneri Richardson: an immuno- and enzyme-histochemical study. J Fish Biol 36, 159–173.

Resseguier, J., Nguyen-Chi, M., Wohlmann, J., Rigaudeau, D., Salinas, I., Oehlers, S.H., Wiegertjes, G.F., Johansen, F.E., Qiao, S.W., Koppang, E.O., Verrier, B., Boudinot, P., Griffiths, G., 2023. Identification of a new mucosal lymphoid organ below the pharynx of teleost fish: tonsils in fish? bioRxiv, 2023.2003.2013.532382.

Romano, N., Taverne-Thiele, J.J., Van Maanen, J.C., Rombout, J., 1997. Leucocyte subpopulations in developing carp (Cyprinus carpioL.): immunocytochemical studies. Fish & Shellfish Immunology 7, 439–453.

Rombout, J.H., Abelli, L., Picchietti, S., Scapigliati, G., Kiron, V., 2011. Teleost intestinal immunology. Fish Shellfish Immunol 31, 616–626.

Ryo, S., Wijdeven, R.H., Tyagi, A., Hermsen, T., Kono, T., Karunasagar, I., Rombout, J.H., Sakai, M., Verburg-van Kemenade, B.M., Savan, R., 2010. Common carp have two subclasses of bonyfish specific antibody IgZ showing differential expression in response to infection. Dev Comp Immunol 34, 1183–1190.

Salinas, I., LaPatra, S.E., Erhardt, E.B., 2015. Nasal vaccination of young rainbow trout (Oncorhynchus mykiss) against infectious hematopoietic necrosis and enteric red mouth disease. Dev Comp Immunol 53, 105–111.

Salinas, I., Zhang, Y.A., Sunyer, J.O., 2011. Mucosal immunoglobulins and B cells of teleost fish. Dev Comp Immunol 35, 1346–1365.

Sepahi, A., Casadei, E., Tacchi, L., Munoz, P., LaPatra, S.E., Salinas, I., 2016. Tissue Microenvironments in the Nasal Epithelium of Rainbow Trout (Oncorhynchus mykiss) Define Two Distinct CD8alpha+ Cell Populations and Establish Regional Immunity. J Immunol 197, 4453–4463.

Sepahi, A., Kraus, A., Casadei, E., Johnston, C.A., Galindo-Villegas, J., Kelly, C., Garcia-Moreno, D., Munoz, P., Mulero, V., Huertas, M., Salinas, I., 2019. Olfactory sensory neurons mediate ultrarapid antiviral immune responses in a TrkA-dependent manner. Proc Natl Acad Sci U S A 116, 12428–12436.

Solem, S.T., Stenvik, J., 2006. Antibody repertoire development in teleosts--a review with emphasis on salmonids and Gadus morhua L. Dev Comp Immunol 30, 57–76.

Sosa, G.A., Roux, M.E., 2004. Development of T lymphocytes in the nasal-associated lymphoid tissue (NALT) from growing Wistar rats. Clin Dev Immunol 11, 29–34.

Tacchi, L., Larragoite, E.T., Munoz, P., Amemiya, C.T., Salinas, I., 2015. African Lungfish Reveal the Evolutionary Origins of Organized Mucosal Lymphoid Tissue in Vertebrates. Curr Biol 25, 2417–2424.

Tacchi, L., Musharrafieh, R., Larragoite, E.T., Crossey, K., Erhardt, E.B., Martin, S.A.M., LaPatra, S.E., Salinas, I., 2014. Nasal immunity is an ancient arm of the mucosal immune system of vertebrates. Nat Commun 5, 5205.

Takizawa, F., Dijkstra, J.M., Kotterba, P., Korytar, T., Kock, H., Kollner, B., Jaureguiberry, B., Nakanishi, T., Fischer, U., 2011. The expression of CD8alpha discriminates distinct T cell subsets in teleost fish. Dev Comp Immunol 35, 752–763.

Tatner, M.F., 1986. The ontogeny of humoral immunity in rainbow trout, Salmo gairdneri. Vet Immunol Immunopathol 12, 93–105.

Tatner, M.F., Manning, M.J., 1983. The ontogeny of cellular immunity in the rainbow trout, Salmo gairdneri Richardson, in relation to the stage of development of the lymphoid organs. Dev Comp Immunol 7, 69–75.

van Poppel, M.N., van den Berg, T.K., van Rees, E.P., Sminia, T., Biewenga, J., 1993. Reticulum cells in the ontogeny of nasal-associated lymphoid tissue (NALT) in the rat. Cell Tissue Res 273, 577–581.

Wosen, J.E., Mukhopadhyay, D., Macaubas, C., Mellins, E.D., 2018. Epithelial MHC Class II Expression and Its Role in Antigen Presentation in the Gastrointestinal and Respiratory Tracts. Front Immunol 9, 2144.

Zapata, A., Diez, B., Cejalvo, T., Gutierrez-de Frias, C., Cortes, A., 2006. Ontogeny of the immune system of fish. Fish Shellfish Immunol 20, 126–136.

Zhang, Y.A., Salinas, I., Li, J., Parra, D., Bjork, S., Xu, Z., LaPatra, S.E., Bartholomew, J., Sunyer, J.O., 2010. IgT, a primitive immunoglobulin class specialized in mucosal immunity. Nat Immunol 11, 827–835.

